# Moving towards precision TMS? Evaluating individual differences and reproducibility of personalized stimulation targets in UK Biobank

**DOI:** 10.1101/2024.04.16.589734

**Authors:** Ying Zhao, Yi-Jie Zhao, Hailun Cui, Richard A.I. Bethlehem, Valerie Voon

**Affiliations:** Institute of Science and Technology for Brain-Inspired Intelligence, Fudan University, Shanghai, China; Department of Psychiatry, University of Cambridge, Cambridge, United Kingdom; Department of Psychology, University of Cambridge, Cambridge, United Kingdom

## Abstract

**Objective:** Personalized transcranial magnetic stimulation (TMS) targeting, guided by functional connectivity (FC), shows potential in treating depression. The present study aims to map individual FC peak location using UK Biobank, to evaluate individual differences and reproducibility of FC-based targets.

**Methods:** We analyzed UK Biobank resting-state fMRI (rfMRI) of 35,423 participants, identifying individual FC peak locations on the dorsolateral prefrontal cortex (DLPFC) that functionally connected to the subcallosal cingulate, amygdala, and ventromedial prefrontal cortex, respectively. Euclidean distance between each participant’s individual peak and group-average peak was calculated. With follow-up rfMRI of 1341 participants, within-subject FC peak location changes were calculated. We also compared common TMS targets and random locations for their median distance to individual peaks in a permutation test.

**Results:** Seed-based FC analyses revealed large differences in the individual FC peak location on DLPFC: the mean distance from the individual peaks to group-average peak ranged from 14.24 to 29.92mm; 70% to 94% of participants were >10mm away from the group-average peak and potentially located outside of the TMS effective area with common TMS coils. Similar variability was observed in within-subject peak locations across two fMRI assessments. Common TMS targets and the group-average FC peak showed no significant difference in median distances to individual FC peaks when compared to random locations.

**Conclusions:** FC peak location shows wide inter- and intra-individual variability. We emphasize a role for individualized TMS neuronavigation targeting but emphasize the need for more reliable biomarker studies.

## Introduction

Depression is a major public health issue associated with marked morbidity, disability and financial burden (1). Up to 30% of patients with depression remain refractory despite adequate conventional interventions such as antidepressants and psychotherapy. Non-invasive neuromodulation techniques such as repetitive transcranial magnetic stimulation (rTMS) show efficacy in the treatment of depression (2). Traditional TMS targets on the brain are mostly based on anatomical landmarks and coordinates, and adopting uniform targets across patients. Advances in neuroimaging have facilitated neuronavigation-guided personalized stimulation based on the individual patient’s brain images, or functional/anatomical connectivity generated from MRI, which have been suggested to potentially show greater clinical efficacy. For example, resting-state functional MRI (rfMRI) functional connectivity (FC) between the left dorsolateral prefrontal cortex (DLPFC) and subcallosal cingulate cortex in depression patients, has suggested potential to guide individualized TMS targeting (3–6).

Systematic reviews of small studies suggest potential utility in personalized neuronavigation targeting compared to standard TMS clinical practice using the 10-20 EEG system F3, Beam F3 or 5.5 cm. Nevertheless, the use of neuronavigation guidance remains to be demonstrated to be more effective than not. A major issue is that any added benefit with neuronavigation guidance may be a small effect size and would require very large sample sizes in randomized controlled trials to demonstrate an effect (7–9). Furthermore, individualized neuronavigation is also less cost-efficient and harder to generalize and scale requiring both a neuronavigator, MRI time, expertise in MRI analysis and additional time and expertise in targeting. An alternative is the use of functional MRI connectivity-based targeting using group-average based rather than individualized targets.

The present study uses large-scale data from over 35,000 individuals in the UK Biobank dataset to explore individual versus group-based functional connectivity of brain regions relevant to depression and rTMS. We compare the between-subject individual peak location differences and group-based peak connectivity of the left DLPFC with the subcallosal cingulate (3–5). The connectivity of the left DLPFC with the amygdala and ventromedial prefrontal cortex (VMPFC) were also analyzed in the same way, given their importance in mood disorders and other psychiatric disorders (10–13). Additionally, we evaluate the stability of the individual FC peak by comparing within-subject individual peaks changes across repeated scans in a subset of over 1,300 individuals who underwent two scans at different times.

## Methods

### Participants

We used UK Biobank (project ID 64044) resting fMRI data for the FC analysis. In the first imaging scan, 35,423 participants passed rfMRI data quality control and were included in the analysis. In addition, 1341 participants with a second time resting fMRI scan were used to investigate within-subject reproducibility of the FC results. Participants demographics are reported in Table S1 and Table S2 in the supplemental material. All participants provided written informed consent.

### Seed-based resting-fMRI functional connectivity

Three regions-of-interest (ROIs), including the subcallosal cingulate, amygdala, and VMPFC were used in seed-based rfMRI FC analysis, respectively. The subcallosal cingulate ROI was defined using the same coordinates (6,16-10) and radius 10 mm as in Fox et al (14, 15) (white matter voxels were excluded). The amygdala ROI were identified from Automated anatomical labelling atlas (AAL3) (16); and VMPFC ROI was identified from the Harvard-Oxford atlas (http://fsl.fmrib.ox.ac.uk/fsl/fslwiki/Atlases).

The preprocessed rfMRI data were provided by the UK Biobank. With preprocessed rfMRI, we further applied spatial smoothing 6mm and regressed out white matter, cerebrospinal fluid, and global mean signal. The FC was calculated in each participant’s native space using ‘y_SCA’ function (17) calculating the Pearson correlation (fisher z-transformed) between time courses of the seed ROI and every other grey matter voxel. Each participant’s FC map was then warped back to MNI space for voxel-wise analysis at group level. Brain results were mapped using Matlab package BrainNet Viewer (18).

### Individual peak and group-average peak in functional connectivity

With each ROI corresponding FC maps, we first calculated the group-average and standard deviation (SD) FC map, respectively. To identify peaks on DLPFC, and given differing means of identifying DLPFC boundaries, we used two methods to define the ‘search area’ as DLPFC: (1) Brodmann area 9 and 46 (BA9+BA46) (19); (2) the left middle frontal gyrus (MFG) from AAL3. Then we threshold the group mean map at FC <-0.1, to identify the group peak location within DLPFC.

In each ‘search area’, we calculated each participant’s FC peak value and identified the peak voxel location (one voxel per participant). The peak voxel value was binarized to 1 and we overlay all participants peak voxels onto one brain to create a heatmap, where voxels with higher values represent a greater number of individual peaks. In addition, we calculated the Euclidean distance between the group-average peak voxel and the individual peak voxel to understand the extent of the distance, specifically to ask if standard TMS coils, e.g. figure-8 coil which can excite cortical area less than 1cm (20–22) in radius cover this distance (23)?

As a subgroup of participants were scanned twice with rfMRI, we compared the peak locations within-subject by calculating the Euclidean distance between the two peak voxels from two time-points.

### Overlay of group peak on DLPFC from different seed regions

To understand how DLPFC connections relate to different seed regions, we overlaid the group peak on DLPFC from each FC analysis. In addition, we overlaid the common TMS target coordinates, including F3 (24), Beam F3, and Rule 5.5cm (25), on the same brain figure to compare their locations with the present study.

### Comparing group-based TMS targets to random locations

We then asked if group-based targets, including the standard TMS targets of F3, beam F3, Rule 5.5cm, and our defined group-average FC peak location, performed better than random locations, in terms of their median distance to 35,423 participants individual peak location. To answer this question, we conducted a permutation test and select 100 random voxels in (1) the BA9+BA46 area and (2) the middle frontal gyrus. For each random voxel, we calculated the Euclidean distance between this voxel and individual peak voxel of 35,423 participants. Lastly, we calculate the median value of the Euclidean distance to represent how good each random voxel might be in its distance to individual peaks. In this way, we build a distribution of 100 random voxels median distance to individual peaks in (1) the BA9+BA46 and (2) the middle frontal gyrus. For the standard TMS targets and the group-average FC peaks identified in the present study, we also calculated their median distance to individual peaks, and identify their rank among the median of 100 random voxels. In this way, we quantify if standard TMS targets or group FC target can statistically perform better than random voxels (at p<0.05). This analysis thus can contribute to the question of whether group-based targets or standard TMS targets are better than random targets in the DLPFC, and if it is necessary to accurately localize a TMS target on the individual brain.

## Results

### Seed-based resting fMRI functional connectivity

Seed-based FC analyses were conducted using each a ROI, namely the subcallosal cingulate, amygdala, and VMPFC at time 1 and time 2 for each individual. The group-average FC mean map and standard deviation are almost identical at time 1 and time 2 (Figure 2a for subcallosal cingulate, Figure 3a for amygdala, and Figure S1 and supplementary results in the online supplement for VMPFC).

**Figure 1.**
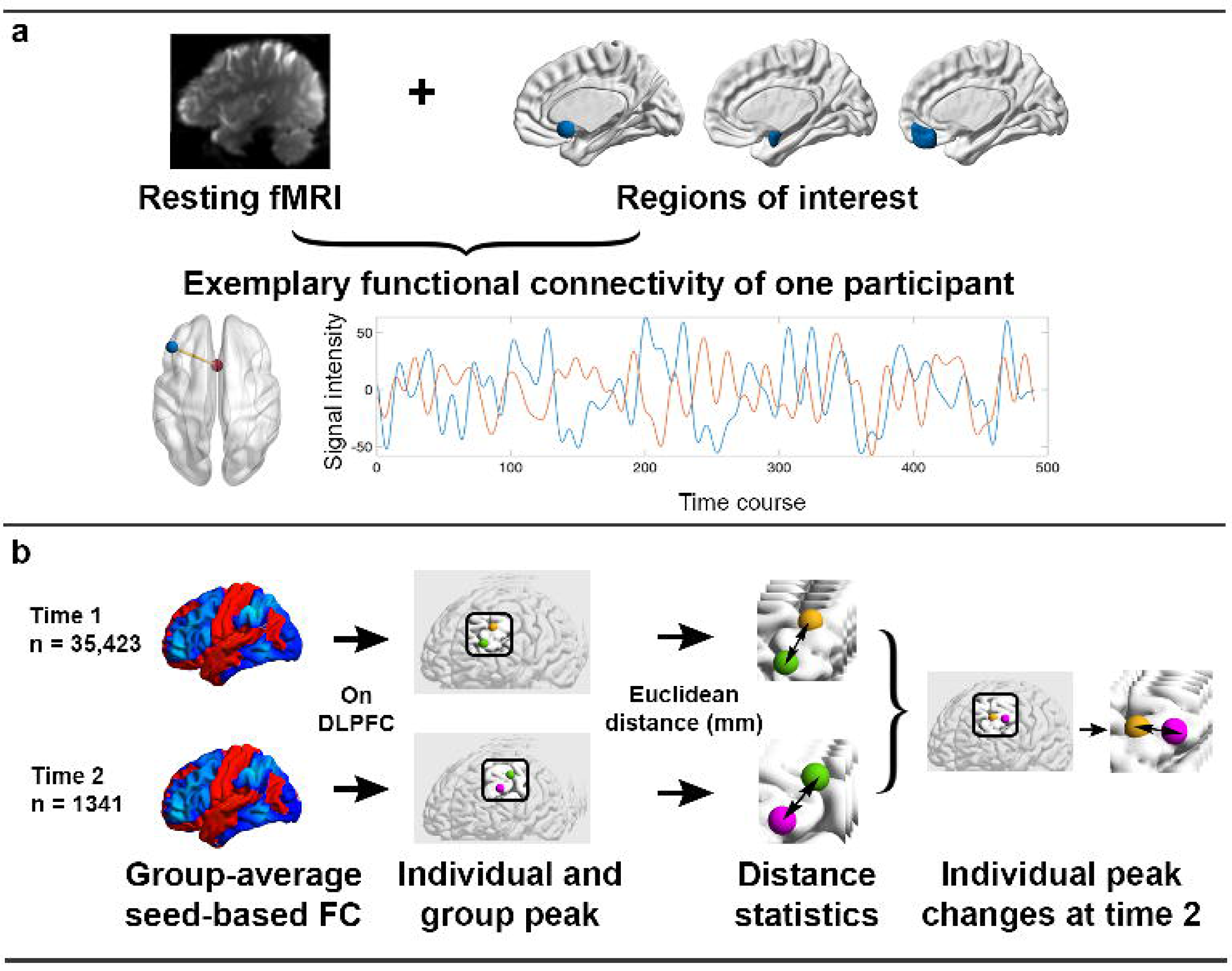
Study overview. a. Seed-based functional connectivity (FC) analysis for one participant. Using the resting fMRI scans, seed-based FC analysis was conducted from three regions of interest (from left to right: the subcallosal cingulate, amygdala, and the ventromedial prefrontal cortex). As illustrated, the individual peak FC on dorsolateral prefrontal cortex (DLPFC) was considered as the personal TMS target. **b. Group level analysis at two time-points.** At each time-point, the group-average FC peak was identified (referred to as the group peak, denoted by the green node). An individual peak voxel was illustrated in orange node at time 1 and the purple node at time 2. Inter-individual difference was quantified by measuring the voxel location distance between individual and group peak in Euclidean distance in millimeter (mm). Furthermore, with repeated resting fMRI scans of 1341 participants, within-subject reproducibility was assessed through calculating peak voxel location changes across two timepoints.

**Figure 2.**
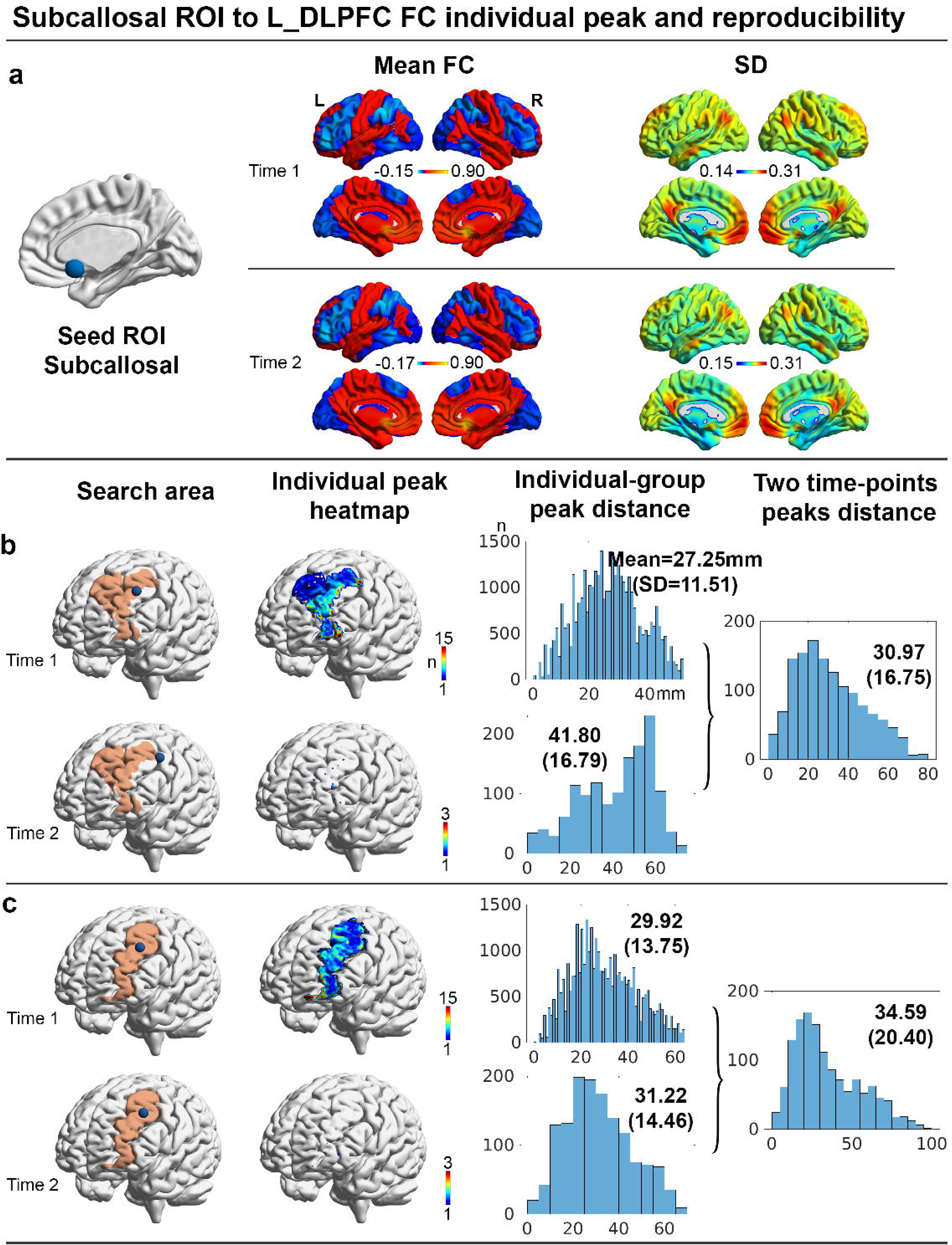
Subcallosal cingulate seed-based functional connectivity. a. Group FC mean and standard deviation. The first and second time-points results are given in two rows and their patterns are similar. **b. Individual peak and its distance to group-average peak on the Broadmann 9 and Broadmann 46 (BA9+BA46). c. Individual peak and its distance to group-average peak on the middle frontal gyrus.** In panel **b** and **c**, the first column brains show the “search area” of DLPFC defined in Broadmann areas or the middle frontal gyrus, respectively. The group-average peak is illustrated in the blue node. The heatmaps show where the individual peaks overlay (one voxel per participant). Higher value in a voxel means more participants’ peaks overlay there. The first histogram shows the Euclidean distance distribution between individual peak and group-average peak. Mean values and standard deviation (SD) are given on the Figure. The second histogram shows the distribution of within-subject peaks in two time-points. ‘n’ denotes participants counts.

**Figure 3.**
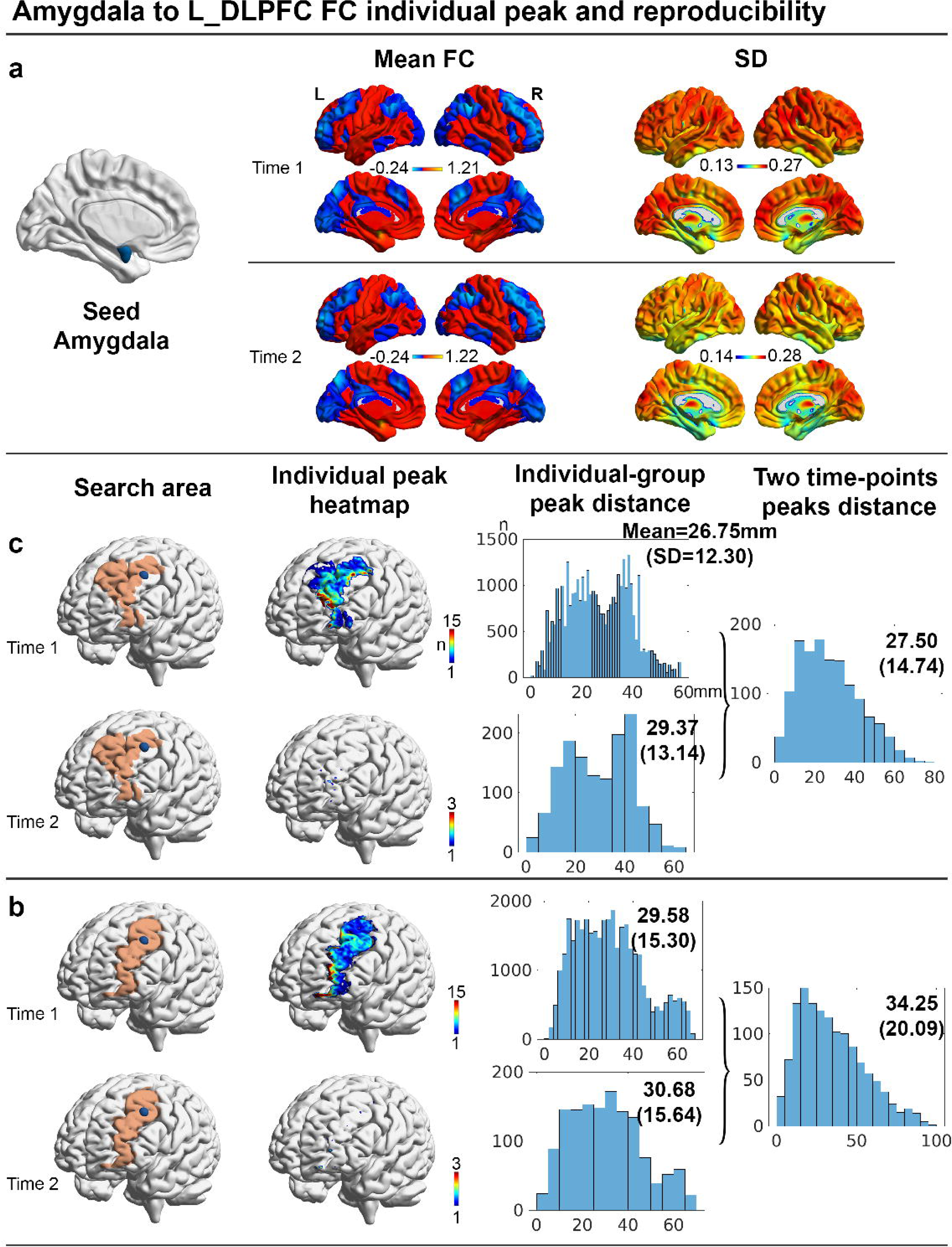
Amygdala seed-based functional connectivity. a. Group FC mean and standard deviation. The first and second time-points results are given in two rows and their patterns are similar. **b. Individual peak and its distance to group-average peak on the Broadmann 9 and Broadmann 46 (BA9+BA46). c. Individual peak and its distance to group-average peak on the middle frontal gyrus.** In panel **b** and **c**, the first column brains show the “search area” of DLPFC defined in Broadmann areas or the middle frontal gyrus, respectively. The group-average peak is illustrated in the blue node. The heatmaps show where the individual peaks overlay (one voxel per participant). Higher value in a voxel means more participants’ peaks overlay there. The first histogram shows the Euclidean distance distribution between individual peak and group-average peak. Mean values and standard deviation (SD) are given on the Figure. The second histogram shows the distribution of within-subject peaks in two time-points. ‘n’ denotes participants counts.

### Subcallosal cingulate

Subcallosal cingulate was negatively connected to a large area on DLPFC (Figure 2a). In the group-average FC map, the peak voxel MNI coordinates are - 44,36,38(the group peak in the “search area” BA9+BA46, Figure 2b), individual peaks can be found with a maximum subjects count=211 (MNI coordinates: −54,10,50). In other words, from more than 35,000 individuals, individual peak voxels were shared amongst 211 subjects. The Euclidean distance between individual and group peak was large with mean=27.25mm (SD=11.51). The number of participants who showed a distance larger than 10mm to the group peak counts was 33,032, which means 93% of the participants are probably out of effective areas of TMS (assuming a 10mm radius) when stimulating at the group peak. The number of participants who showed a distance larger than 20mm to the group peak counts was 25,341, which means 72% of the participants are probably out of effective areas of TMS if assuming a 20mm radius. Similarly within-subject reproducibility showed wide variability: the two time-points within-subject peaks distance was similarly large with mean=30.97mm (SD=16.75).

In MFG (Figure 2c), individual peaks can be found with a maximum subjects count=143 (MNI coordinates: −26,−4,58). In other words, from more than 35,000 individuals, individual peak voxels were shared amongst 143 subjects. The Euclidean distance between individual and group peak was large with mean=29.92mm (SD=13.75). The number of participants who showed a distance larger than 10mm to the group peak counts was 33,425, which means 94% of the participants are probably out of effective areas of TMS (assuming a 10mm radius) when stimulating at the group peak. The number of participants who showed a distance larger than 20mm to the group peak counts was 26,204, which means 74% of the participants are probably out of effective areas of TMS if assuming a 20mm radius. Similarly, within-subject reproducibility showed wide variability: the two time-points within-subject peaks distance was similarly large with mean=34.59mm (SD=20.40).

### Amygdala

The amygdala was negatively connected to a large area on DLPFC (group-average FC peak at MNI coordinates −42,30,40, Figure 3a). In the “search area” BA9+BA46 (Figure 3b), individual voxel peaks can be found shared with a maximum subjects count=132 (MNI coordinates: −36,58,12). The Euclidean distance between individual and group peak was similarly large with a mean=26.75mm (SD=12.30). The number of participants who had a distance larger than 10mm to the group peak counts=32,311, which means 91% of the participants are likely out of effective areas of TMS when stimulating group-average peak (assuming a 10mm radius). The number of participants who showed a distance larger than 20mm to the group peak counts was 23,479, which means 66% of the participants are probably out of effective areas of TMS if assuming a 20mm radius. Reproducibility was similarly broad with the two time-points within-subject peaks distance showing large mean=27.50mm (SD=14.74).

In the “search area” MFG (Figure 3c), individual voxel peaks can be found shared with a maximum subjects count=127 (MNI coordinates: −32,60,22). The Euclidean distance between individual and group peak was similarly large with a mean=29.58mm (SD=15.30). The number of participants who had a distance larger than 10mm to the group peak counts=32,414, which means 92% of the participants are likely out of effective areas of TMS when stimulating group-average peak (assuming a 10mm radius). The number of participants who showed a distance larger than 20mm to the group peak counts was 24,248, which means 68% of the participants are probably out of effective areas of TMS if assuming a 20mm radius. Reproducibility was similarly broad with the two time-points within-subject peaks distance showing large mean=34.25mm (SD=20.09).

### Overlay of group peak on DLPFC from different seed regions

The overlap of group peak on DLPFC areas from the subcallosal cingulate, amygdala, and VMPFC are shown in Figure 4 (Figure 4a shows group peak within BA9+BA46 and Figure 4b shows within MFG). The three common clinically used TMS targets coordinates, including F3, beam F3, and Rule 5.5cm are mapped on the same brain (Figure 4) (14). The six nodes are close to each other on the rostral middle frontal gyrus. The F3 and 5.5cm rule target is the furthest, with 28mm apart (Figure 4a within BA9+BA46). Given the nodes locations and distance, it is possible to cover all these nodes with a coil that can excite cortical area more than 14mm in radius.

**Figure 4.**
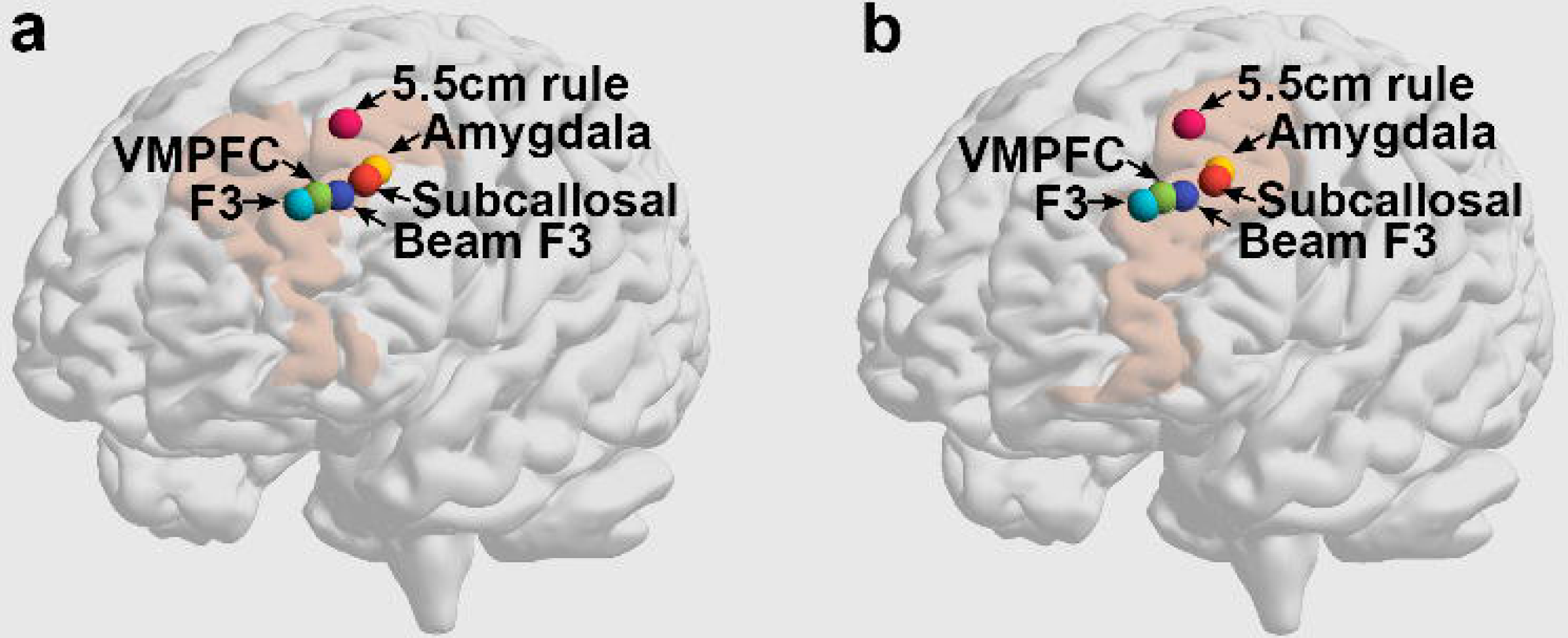
Overlay of group FC peak and common TMS targets. The figure illustrates where group FC peak from the subcallosal, amygdala, and VMPFC locates in the Brodmann area 9 and 46 (the left brain) or in the middle frontal gyrus (the right brain). All nodes locate within middle frontal gyrus regardless DLPFC definition. The F3 and 5.5cm rule target is the furthest, with 28mm apart. Given the nodes locations and distance, it is possible to cover all these nodes with a coil that can excite cortical area more than 14mm in radius.

### Comparing group-based TMS targets to random locations

The median distance between group-based TMS targets and individual FC peaks were not significantly better than random voxels (non-significant at p<0.05). In the permutation test, with the subcallosal cingulate seed FC analysis, we calculated the Euclidean distance between the group FC peak to 35,423 participants’ individual peaks. We also calculated the same distance measure for common TMS targets, namely F3, beam F3, Rule 5.5cm. In addition, we calculated the same distance measure for 100 random voxels. The ranks of median values of the random voxels and group-based targets is given in Figure 5a within BA9+BA46 or in Figure 5b within MFG areas. None of the common TMS targets or group FC peak performed better than random locations (non-significant at p<0.05).

**Figure 5.**
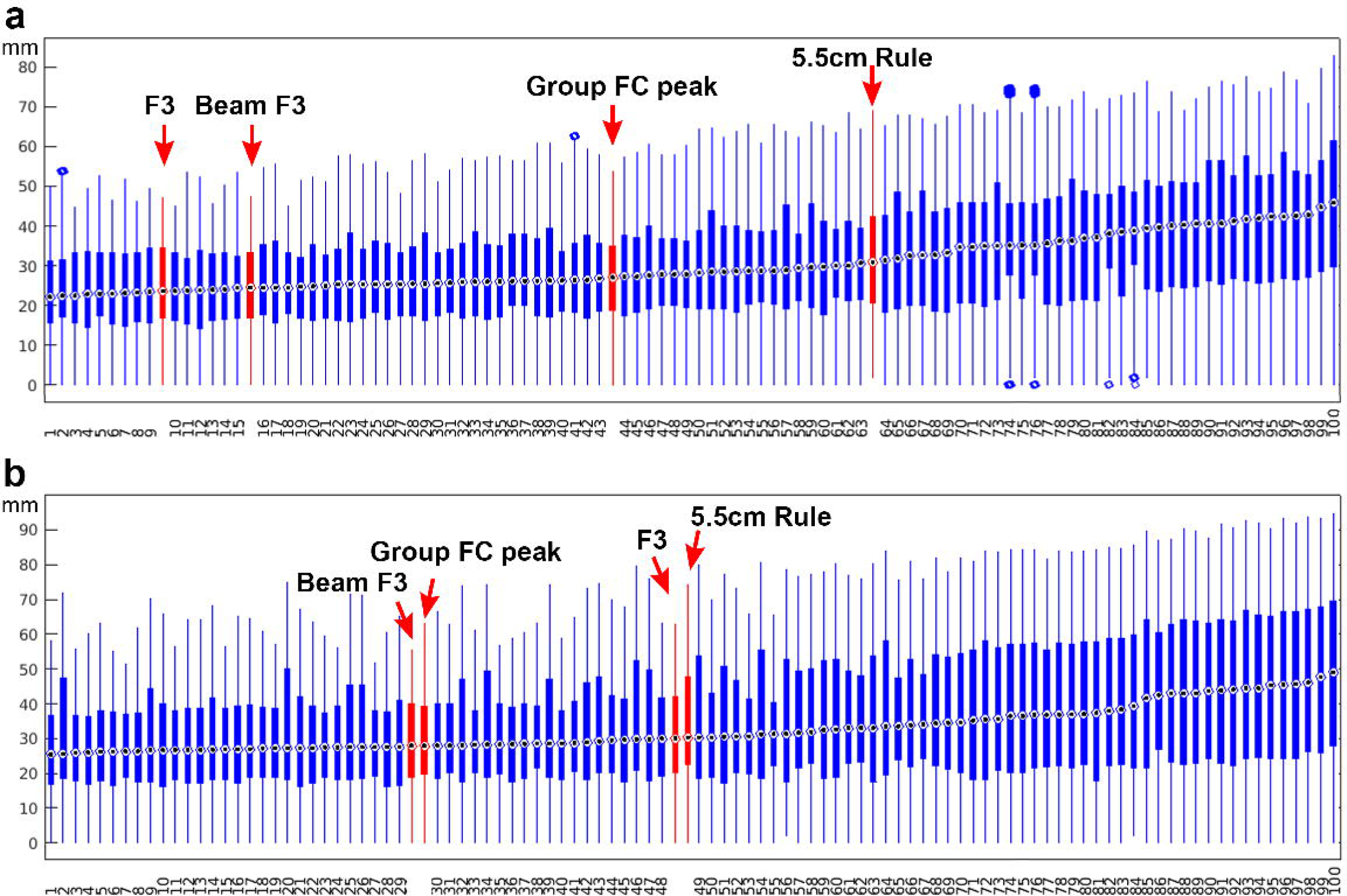
Boxplot showing Euclidean distance of the common TMS targets and random voxels to individual FC peaks. a. DLPFC defined in the Broadmann area 9 and 46. b. DLPFC defined in the middle frontal gyrus. The seed region is the subcallosal cingulate. Each box shows the distance between a voxel and 35,423 participants’ individual FC peak median value (the central dot), 25th, and 75th percentiles, and the whiskers extend to the minimum and maximum values. The Blue circles denote the outlier values. Neither the group FC peak nor the common TMS targets could perform better than top 5 random voxels, indicating they were not performing better than random voxels (non-significant at p<0.05).

## Discussion

Using UK Biobank resting-state fMRI of 35,423 participants and repeated resting-state scans of 1341 participants, our studies highlight two critical findings in the question of rTMS targeting of the DLPFC for depression treatment focusing on anticorrelation (negative FC) of the DLPFC and subcallosal cingulate. Firstly, the inter-individual variability in FC is substantial: the majority of participants’ peak FC fell out of the 10mm radius if we use group-average coordinates as the target. Similarly, the performance of standard TMS targeting of F3, Beam F3 or the 5.5 cm rule along with the group peak FC performed no better than random targeting within the DLPFC in terms of relationship to individual peaks. Secondly, the repeated resting-state fMRI scan showed large within-participants variance of peak locations although the group peak was the same or very close between scans, which raises a further question of the stability of these individual peak locations obtained from scans at different time points.

### Functional connectivity of individual differences

In the current study, we explored individual differences in FC peak locations that connected to subcortical ROIs. Smaller pilot studies suggest potentially greater efficacy following FC guided personalized TMS targeting of DLPFC-subcallosal cingulate hypoconnectivity for depression treatment, albeit with a small effect size (3, 5). However, an iTBS study comparing anatomical-based neuronavigation (border anterior and middle third of MFG) and standard TMS targeting without neuronavigation in 37 depressed inpatients was halted early due to lack of difference in outcomes(7). A recent large iTBS study of FC neuronavigation guided targeting in the DLPFC to the anterior insula compared to without neuronavigation targeting F3 also did not show any differences in antidepressant efficacy (9). The Stanford Neuromodulation Therapy (SNT) protocol which uses high dose accelerated neuronavigation-guided targeting shows marked efficacy but whether this is related to the accelerated design or the higher doses as compared to neuronavigation remains to be established. Further largescale FC-guided neuronavigation studies are underway.

An alternate approach is to consider targeting group FC peaks. Indeed, we show that the group FC is remarkably stable over time with similar group FC in 1341 participants across a 2-year interval. However, we found significant differences between group peak and individual peak coordinates across different ROIs (including the subcallosal cingulate, amygdala, and VMPFC), with a large proportion of subjects falling out of the 10mm range of group peak locations, which is generally recognized as the effective coverage of TMS. Thus, assuming that individual peak FC of DLPFC to subcallosal cingulate might have therapeutic relevance, our data suggests that using group FC peaks with standard Figure-of-8 coils with 10mm radius effect, or even with larger coils covering 20mm radius, would have limited utility in coverage of individual FC peaks.

Measuring correlations between brain regions based on the ongoing fluctuations of the BOLD signal are subject to various neural and non-neural confounding factors (26). Apart from the MRI mechanistic and technical causes, inter-subject variability in functional connectivity is inherently connected to brain evolution and development, with evidence revealing the highest variability in phylogenetically late-developing regions like the frontal, temporal, and parietal association cortex areas, suggesting a role in evolutionary cortical expansion (27). Although difficulties exist in establishing a universal target in TMS treatment, these distinctive patterns of FC can serve as a biomarker for parcellating heterogeneity of disorders and has been considered as elements of personalized, precision medicine (28–30).

### Functional connectivity within-subject reproducibility

Critically, the within-subject individual FC across two time points also showed a large difference, an effect potentially greater than the individual-group peak distance, highlighting the low stability and reproducibility of the personal FC peak in rfMRI. A meta-analysis consolidating existing knowledge on test-retest reliability of edge-level functional connectivity found a low intraclass correlation coefficient (ICC) of 0.29, indicating marked instability among studies despite adequate artifact removal methods (31). Using data from the Human Connectome Project, Ning et al. showed that, using repetitive MRI scan settings involving two 15-minute scans on consecutive days with a 121Zmm FWHM smoothing kernel, the intra-subject variability of subcallosal cingulate-DLPFC functional connectivity maps was 26.31Zmm, whereas a single 15-minute scan led to a larger inter-scan distance of 37.41Zmm (32). Our result echoed the conclusion derived from Human Connectome Project data and have expanded the intra-subject variability. Even though technical developments might allow higher resolution in fMRI (e.g. using 7T MRI) and higher signal-to-noise ratio with improved data processing parameters may allow reproducible individualized targets in the future, the low within-subject stability and large individual differences might have a basis not in MRI techniques but in the dynamic nature of brain resting-state, e.g. dynamic FC (33) and thus the peak location changes. Therefore, our findings highlight marked inter- and intra-individual variability in DLPFC and subcallosal cingulate FC targeting. The question remains whether personalization based on individual FC is technically feasible for reliably predicting effective targets and if the increased precision, entailing both human and technical costs, can significantly outperform the clinical efficacy achieved by employing conventional targeting methods.

### Alternative solutions

We further highlight that using group FC targeting or the standard TMS targets using F3, Beam F3 or the 5.5 cm rule performs no differently from random FC targets within the DLPFC in its relationship to individual FC targets. Our findings suggest likely similar efficacy between these common TMS targets assuming the relevance of DLPFC and subcallosal FC as a biomarker for rTMS antidepressant treatment efficacy. Further large scale randomized controlled trials assessing this personalized neuronavigation targeting are eagerly awaited.

Targeting the left DLPFC with rTMS is clearly effective for refractory depression. Therapeutic outcome remains limited at around 40% clinical response rate for conventional approaches (34). Identifying biomarker targets to optimize therapeutic outcomes is critical and can include either optimizing targeting approaches such as (i) greater precision in targeting within the DLPFC, alternative approaches involving other forms of physiology such as simultaneous collection of galvanic skin responses; (ii) targeting alternate networks including the dorsomedial prefrontal cortex or orbitofrontal cortex; (iii) identifying heterogeneity in major depression that might be responsive to different targets; or optimizing delivery approaches such as accelerated designs or higher doses.

Our findings highlight a role for personalized neuronavigation targeting in clinical management. However, its necessity may depend on the different needs of the population and goals. Neuronavigation and precision targeting is clearly necessary in an academic research context. However, neuronavigation is costly both in terms of need of a MRI, specialized personnel, equipment and time, hence perhaps limiting its clinical utility particularly at scale in the general population in community settings and local hospitals or deprived areas with limited funds, resources, personnel and high patient needs. Optimal TMS treatment in specialized centers may benefit from individualized targeting.

Given the limitations of the group-based targets and reproducibility issue in FC-guided personalized targets, another plausible solution might be the use of coils with larger effective coverage with lower focality (35), e. g. the H-coil and double-cone coil, that cover a larger region on the DLPFC to ensure a higher likelihood of individual peaks being simulated (36); likewise, broad spectrum antibiotics can work on a large range of bacteria which do not necessarily require personalized treatment. A limited number of studies have compared the response rate amongst such coils. For example, the H-coil showed a 46% response rate at four weeks of treatment which increased to 81% at 22 weeks; double-cone coils have reported a response rate of 40-52.4% (37). One review reported noticeable higher response rate with H-coil than figure-8 coil (38) while a randomized controlled trial reported no significant difference in response rate between the double-cone coil and figure-8 coil (39). Further clinical trials may be indicated to determine if these are necessarily more effective than standard coils.

Given how common depression and/or psychiatric disorders occur at the population level, the financial and time burdens for patients and national health services (e.g. NHS in UK and developing economies) are not negligible (40), brain stimulation covering larger areas on the DLPFC might be a more feasible route towards treatment. When selecting different coils, the heatmap (e.g. in Figure 2, 3, and S1) of individual peak voxels based on population dataset could be used to guide which coil to consider. When determining clinical interventions for an individual, a staged treatment plan might be indicated with coils targeting broader regions first, then personalized treatment requiring greater financial and time costs indicated for severe or non-responsive patients.

### Limitations

Our findings are not without limitations. The time gap between the two rfMRI scanning is an average of 2.3 years. Therefore, the stability of individual peak (within-subject) in the context of different time gaps, e.g. days and weeks, require further investigation. However, reproducibility studies described above further highlight within-individual variability.

In conclusion, potential brain stimulation targets based on functional connectivity, particularly of left DLPFC-subcallosal anticorrelation, show wide inter-individual and temporal variability. We emphasize the need for further studies to optimize biomarker identification for neuronavigation targeting. Further optimization of rTMS clinical efficacy might focus on alternate or combination biomarkers, accelerated designs, inclusion of peripheral physiology, higher doses or targeting alternate networks.

## Acknowledgments

This work is supported by Medical Research Council Senior Clinical Fellowship (Grant No. MR/W020408/1 and No. MR/P008747/1) and the Young Scientists Fund of the National Natural Science Foundation of China (82201670).

