## supplemental material for "Moving towards precision TMS? Evaluating individual differences and reproducibility of personalized stimulation targets in UK Biobank"

**Table S1. Demographics**

|  | **Mean** | **SD** | **Available participates (n)** |
| --- | --- | --- | --- |
| **BMI** | 26.38 | 4.27 | 30,709 |
| **AUDIT** | 5.21 | 4.16 | 21,983 |
| **RDS-4** | 5.20 | 1.75 | 30,068 |
| **PHQ-9** | 2.55 | 3.47 | 21,833 |
| **GAD-7** | 2.00 | 3.22 | 21,918 |

AUDIT: Alcohol Use Disorders Identification Test.

RDS-4: Recent Depressive Symptoms.

PHQ-9: Patient Health Questionnaire, the depression module.

GAD-7: Generalized Anxiety Disorder Assessment.

SD: standard deviation.

**Table S2. Participants statistics in analyses**

|  | **Available participates (n)** | **Age mean (SD)** | **Gender (female percentage)** |
| --- | --- | --- | --- |
| **Time 1** | 35,423 | 63.43 (7.51) | 54% |
| **Time 2** | 1,341 | 64.76 (7.17) | 51% |

SD: standard deviation.

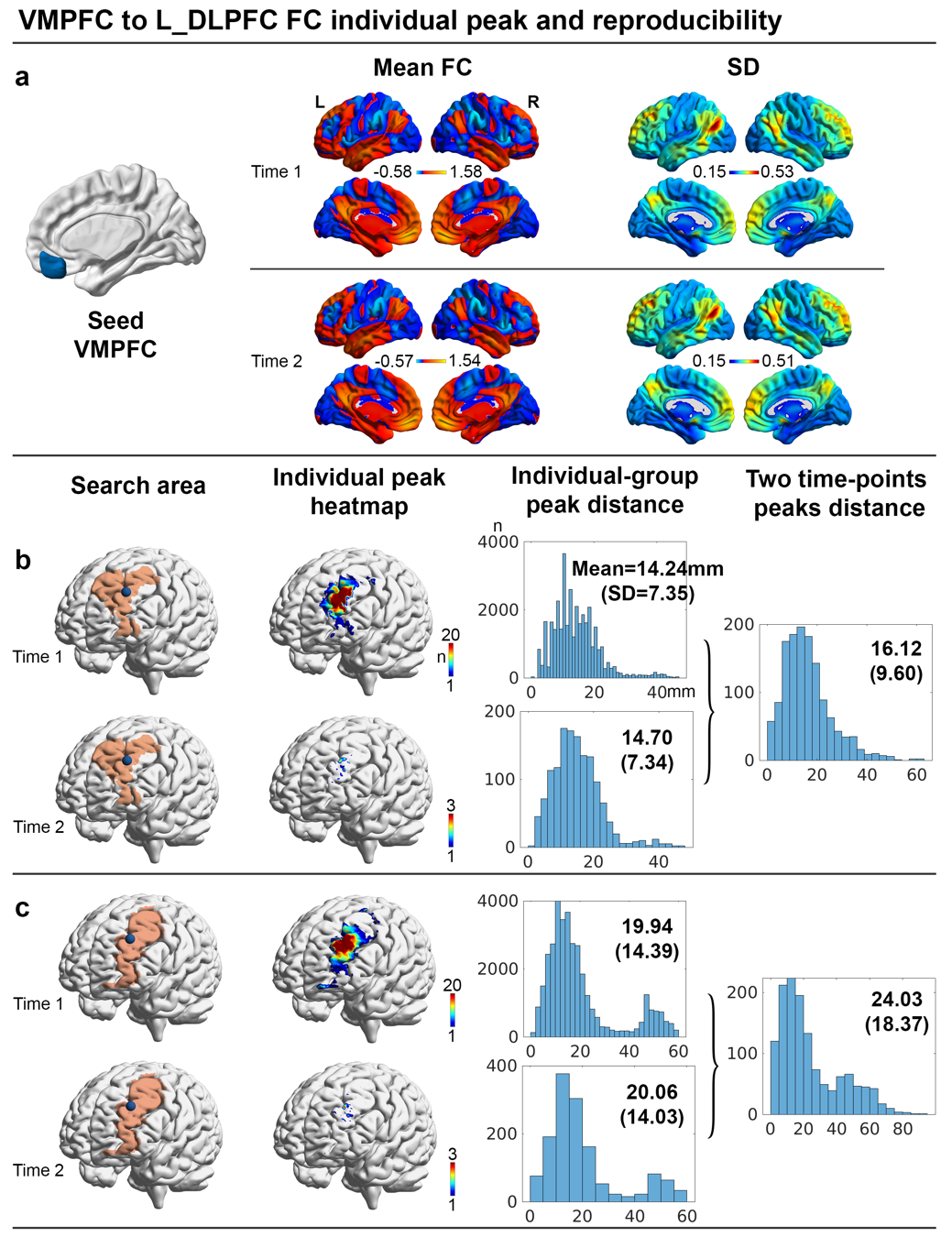

**Figure S1. VMPFC seed-based functional connectivity. a. Group FC mean and standard deviation.** The first and second time-points results are given in two rows and their patterns are similar. **b. Individual peak and its distance to group-average peak on the Broadmann 9 and Broadmann 46 (BA9+BA46). c. Individual peak and its distance to group-average peak on the middle frontal gyrus.** In panel **b** and **c**, the first column brains show the “search area” of DLPFC defined in Broadmann areas or the middle frontal gyrus, respectively. The group-average peak is illustrated in the blue node. The heatmaps show where the individual peaks overlay (one voxel per participant). Higher value in a voxel means more participants’ peaks overlay there. The first histogram shows the Euclidean distance distribution between individual peak and group-average peak. Mean values and standard deviation (SD) are given on the Figure. The second histogram shows the distribution of within-subject peaks in two time-points. ‘n’ denotes participants counts.

**Supplementary results**

**VMPFC Seed-based resting fMRI functional connectivity**

VMPFC was negatively connected to a focal area on DLPFC (Figure S1a). In the group-average FC map, the peak voxel MNI coordinates are -38,46,34. In the “search area” BA9+BA46 (Figure S1b), individual peak can be found with a maximum subjects count=206 (MNI coordinates: -46,44,32). The Euclidean distance between individual and group peak was mean=14.24mm (SD=7.35). The number of participants who had a distance larger than 10mm to the group peak counts=24,819, which means 70% of the participants are probably out of effective areas of TMS (assuming a 10mm radius) when stimulating at the group peak. The number of participants who showed a distance larger than 20mm to the group peak counts was 6232, which means 18% of the participants are probably out of effective areas of TMS if assuming a 20mm radius. Regarding reproducibility, the two time-points within-subject peaks distance was mean=16.12mm (SD=9.60).

In the “search area” MFG (Figure S1c), individual peak can be found with a maximum subjects count=139 (MNI coordinates: -34,50,34). The Euclidean distance between individual and group peak was mean=19.94mm (SD=14.39). Regarding reproducibility, the two time-points within-subject peaks distance was mean=24.03mm (SD=18.37). The number of participants who had a distance larger than 10mm to the group peak counts=27,452, which means 78% of the participants are probably out of effective areas of TMS (assuming a 10mm radius) when stimulating at the group peak. The number of participants who showed a distance larger than 20mm to the group peak counts was 11,123, which means 31% of the participants are probably out of effective areas of TMS if assuming a 20mm radius.
